# Dynamic analysis of lower leg muscles response to whole body vibration stimulation at different frequencies and postures: implications for training

**DOI:** 10.1101/2021.08.31.458312

**Authors:** Isotta Rigoni, Tecla Bonci, Paolo Bifulco, Antonio Fratini

**Affiliations:** Biomedical Engineering, College of Engineering and Physical Sciences, Aston University, Birmingham, United Kingdom; EEG and Epilepsy Unit, Clinical Neuroscience Department, University Hospital and Faculty of Medicine of Geneva, Geneva, Switzerland; Department of Mechanical Engineering and the Insigneo Institute for in silico Medicine, University of Sheffield, Sheffield, UK; Department of Electrical Engineering and Information Technology, University of Naples Federico II, Via Claudio 21, 80125, Naples, Italy

**Keywords:** electromyography (EMG), soft-tissue acceleration, whole body vibration (WBV), frequency, posture

## Abstract

**Purpose:** To characterise the mechanical and neuromuscular response of lower limb muscles in subjects undergoing Whole Body Vibration (WBV) at different frequencies while holding two static postures.

**Methods:** Twenty-five participants underwent WBV at 15, 20, 25 and 30 Hz while holding a static ‘hack squat’ and on ‘fore feet’ posture. Surface electromyography (sEMG) and soft tissue accelerations were collected from Gastrocnemius Lateralis (GL), Soleus (SOL) and Tibialis Anterior (TA) muscles.

**Results:** Only specific WBV settings led to a significant increase in muscle contraction. Specifically, the WBV-induced activation of SOL and GL was maximal in fore-feet and in response to higher frequencies. Estimated displacement at muscle bellies revealed a resonant pattern never highlighted before. After stimulation starts, muscle oscillation reaches a peak followed by a drop and a further stabilisation (few seconds after the peak) that suggests the occurrence of a neuromuscular activation to reduce the vibration-induced oscillation.

**Conclusion:** Lower leg muscles need a response time to tune to a vibratory stimulation, which discourages the use of dynamic exercises on vibrating platforms. To maximize calf muscle response to WBVs, a stimulation frequency in the range of 25-30 Hz and an ‘on fore feet’ posture are recommended.

## Introduction

Whole Body Vibration (WBV) refers to the use of mechanical stimulation, in the form of vibratory oscillations extended to the whole body, to elicit neuromuscular responses in multiple muscle groups. Vibrations are generally delivered through lower limbs via the use of platforms on which subjects stand. Training and rehabilitation programmes utilising this modality often include physical exercise performed on such platforms (Dolny and Reyes, 2008). This approach has become increasingly popular as it evokes a large muscle response and, more importantly, it elicits muscles activity through physiological pathways (Granit and Steg, 1956), improving overall motor performance while enhancing strength and flexibility (Alam et al., 2018; Delecluse et al., 2003; Osawa et al., 2013; Rittweger, 2010; Saquetto et al., 2015; Wyon et al., 2010).

The mechanism believed to be responsible for such outcomes, known as Tonic Vibration Reflex (TVR), has been proven to explain the increased and synchronised motor-unit (MU) firing rates recorded during locally-applied (i.e., focal) vibrations (Burke et al., 1976; Burke and Schiller, 1976). Indeed, when vibrations are applied directly to tendons or muscle bellies, muscle fibres length changes activating a reflex response from muscle spindles. This translates in an increased MU firing rates phased-locked specifically to the vibratory cycle, i.e. the TVR (Burke and Schiller, 1976; Hagbarth et al., 1976; Person and Kozhina, 1992).

In WBV, instead, vibrations are not applied locally, but transferred to the target muscles via the kinematic chain determined by the body posture (Cardinale and Lim, 2003; Pollock et al., 2012; Ritzmann et al., 2010); this provides similar muscular outcomes with respect to focal stimulations as well as additional systemic postural responses, allowing better flexibility and applicability to large exercise programmes (Rittweger, 2010).

Specifically, when the whole body is exposed to mechanical shocks (such as vibrations), absorption strategies act to dampen oscillations and dissipate energy through modulation of both muscle activity and joint kinematics, over which the body has prompt control (Gross and Nelson, 1988; Lafortune et al., 1996). Moreover, in WBV, somatosensory feedback pathways are enhanced by reflexes arising from mechanoreceptors in the lower limbs, with significant implications for motor coordination and postural control during quiet stance (Fitzpatrick et al., 1992). Human ability to keep an upright stance, in fact, depends on the integration of somatosensory, vestibular and visual feedbacks (Horak et al., 1997; Peterka, 2002) that translates in specific muscles activation patterns and body sway about the ankle joint (Winter, 1995). The ankle strategy - actuated by modulation of plantar- and dorsi-flexor activation - is indeed considered the primary approach to reposition the centre of gravity either during quiet stance or in response to external perturbations (Gatev et al., 1999; Winter, 1995). Cross-correlation analyses between muscle activity and centre of pressure further confirm how leg muscles (with particular focus on the Gastrocnemius Lateralis and Soleus) play a central role in postural control dynamics (Fitzpatrick et al., 1992; Gatev et al., 1999). Therefore, it could be inferred that, if selectively targeting lower leg muscles, WBV training may also improve coordination and balance of different subpopulations.

Promising results of WBV training are reported in the literature (Bautmans et al., 2005; Bogaerts et al., 2007; Lam et al., 2012; Mahieu et al., 2006; Ritzmann et al., 2014; Rogan et al., 2011; Torvinen et al., 2002a; Turbanski et al., 2005; van Nes et al., 2004), but a few discording results still jeopardize the systematic use of such approach in training and rehabilitation practices (Torvinen et al., 2003, 2002b; Yang et al., 2015). Conflicting results might be related to the high amount of variability in WBV settings (e.g., stimulation frequency, posture, stimulation amplitude, stimulation duration etc.) used throughout different studies, while still lacking of standardised training protocols. Among the most investigated variables, stimulation frequency and subject posture have relevant impact in eliciting an efficient muscle tuning response to WBV (Lienhard et al., 2014; Pollock et al., 2010; Ritzmann et al., 2013).

Previous findings suggest that muscles contract to reduce the soft-tissue resonance, especially when the stimulation frequency, *ω*_*a*_, is close to their natural one(Wakeling et al., 2003, 2002a; Wakeling and Nigg, 2001a). This process, known as muscle tuning, is perpetrated by muscles to minimize the soft-tissue vibrations (Wakeling and Nigg, 2001b)(Wakeling et al., 2001) and has been recently proposed as one of the possible body reactions to WBV (Abercromby et al., 2007a, 2007b). Therefore, a careful selection of stimulation frequency, *ω*_*a*_, to match the resonant one, *ω*_*0*_, seems the key element to maximise muscle responses to WBVs (Cesarelli et al., 2010). Generally, the natural frequency of a system depends on its mass, *m*, and stiffness, *k*, according to the equation 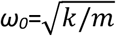 (Rittweger, 2010); being the mass of a muscle constant, modulating muscle stiffness via a change in subject posture results in a change in the system’s natural frequency. This seems to explain conflicting results in the literature when combined dynamic and static exercises (holding a fixed posture e.g. hack squat) are performed on platforms (Bosco et al., 1999; Fagnani et al., 2006) (Wyon et al., 2010). During dynamic exercises on a vibrating platform the body kinematic chain involved in the transmission of the mechanical stimulus changes continuously, making extremely difficult to define the stimulus delivered at the target muscle group. However, during static exercises on a platform, the energy dissipated through joint kinematics is constant and muscle contraction is the major mechanism tuned to dampen vibration. Indeed, Abercromby et al. confirmed that static exercises during WBV enhance higher muscle response than performing dynamic exercises, during which muscles contract in an eccentric and concentric fashion (Abercromby et al., 2007a).

With the present study, the authors want to provide a comprehensive analysis of the effects of WBV on lower leg muscles. In particular, this paper reports the findings on muscle activity and soft-tissue mechanical dynamics and their link to both vibration frequency and subject posture, when WBV is delivered via a side alternating platform. Suggestions are presented on the appropriate approach to use for training professionals and practitioners, with consideration given to potential implications for postural control enhancement training.

## Materials and Methods

### Participants and experimental design

Seventeen females and eight males (age: 24.8 ± 3.4 years; height: 172.0 ± 8.6 cm; mass: 64.6 ± 10.5 kg) volunteered in the study. History of neuromuscular or balance disorders as well as recent injuries were among the exclusion criteria. To evaluate muscle activation and displacement during WBV, surface electromyography (sEMG) signals and accelerations were collected from three lower limb muscles during two static exercises performed in static conditions (without WBV – hereafter called baseline activity) and when different vibration frequencies were delivered.

Pairs of Ag/AgCl surface electrodes (Arbo Solid Gel, KendallTM, CovidienTM 30 mm x 24 mm, centre-to-centre distance 24 mm) were placed over the Gastrocnemius Lateralis (GL), Tibialis Anterior (TA) and Soleus (SOL) muscles of the dominant leg accordingly to SENIAM guidelines (Hermens et al., 2000). The reference electrode was placed on the styloid process of the right ulna. The EMG data were sampled at 1000 Hz (PocketEMG, BTS Bioengineering, Milano, Italy) and sent wirelessly to a laptop via the Myolab software, version 2.12.129.0 (BTS Bioengineering, Milano, Italy).

Accelerations were measured via tri-axial accelerometers (AX3, Axivity Ltd, Newcastle, United Kingdom; range = ±16g, sampling frequency = 1600 Hz) placed on GL, TA and SOL muscle bellies, next to the EMG electrodes. The accelerometers were aligned with the x-axis parallel to the long axis of the leg segment, the z-axis normal to the skin surface and the y-axis perpendicular to the x-y plane. Accelerations were recorded using the open source software OMGUI developed by Newcastle University (“AX3 OMGUI Configuration and Analysis Tool, v38, GitHub,” 2015).

### Whole Body Vibration stimulation protocol

Subjects underwent the WBVs barefoot. The WBVs were delivered via a side-alternating platform (Galileo^®^ Med, Novotec GmbH, Pforzheim, Germany), as it was shown to evoke bigger neuromuscular activations than synchronous vibrating ones (Ritzmann et al., 2013): a peak-to-peak amplitude of 4 mm was used. For each subject, ten trials were collected to evaluate the effect of different stimulation frequencies -0, 15, 20, 25, 30 Hz- and two subject postures: hack squat (HS) and fore-foot (FF). During HS trials, subjects were asked to keep their knees flexed at about 70°, with 0° corresponding to the knee fully extended, and a goniometer was used to check the angle at the beginning of each HS trial. During FF trials, subjects were asked to keep their heels in contact with a parallelepiped-shaped foam (30 × 4 × 3 cm) glued on the platform, to ensure their heels were 3 cm off the ground. Trials were administered in a random order with a one-minute break between consecutive trials.

Hereafter, trials with vibratory stimulation are referred to as “the WBV trials” *(HS*_15_ *HS*_20_ *HS*_25_ *HS*_30_ *FF* _15_*FF*_20_ *FF*_25_ *FF*_30_) and the others as “the baseline trials” (*HS*_0_ and *FF*_0_). WBV trials consisted of 40 seconds: recordings contained 10 seconds with no vibration (*WBV*_*off*_ portion), once the subject acquired the prescribed posture, followed by 30 seconds of WBVs at the prescribed frequency (*WBV*_*on*_ portion). Baseline trials were used to assess the relevant subject-specific EMG baseline activity (*HS*_0_ and *FF*_0_) over a 30 s period.

Twelve Vicon Vero v2.2 optical cameras (Vicon Nexus, Vicon Motion Systems Limited, Oxford, UK) were used to measure subjects’ posture and assure consistency throughout the experiment. Sixteen retroreflective markers were attached to the participant’s body, according to the Plug-In-Gait Lower-Limb model. Data were sampled at 100 Hz and knee and ankle angles were obtained by extracting the kinematics in the sagittal plane using the proprietary software.

### Data processing and features extraction-EMG Data

To isolate the muscle activity preceding the stimulation (*WBV*_*off*_) from the one actually induced by the vibrations (*WBV*_*on*_), each WBV trial was split into two epochs: 10 and 30 seconds, respectively. The central portions of these signals (6 and 20 seconds, respectively) were extracted and retained for analyses. Similarly, the central 20 seconds of the baseline trials (*HS*_0_ and *FF*_0_) were extracted and retained for analyses. In total, ten 20 second-long epochs and eight 6 second-long epochs were analysed for each muscle and subject.

All epochs were band-pass filtered between 5 and 450 Hz with a 5^th^ order Butterworth filter and a mean running root mean square (*rRMS*) value was obtained from both the baseline (*RMS*_*baseline*_) and the *WBV*_*off*_ epochs (*RMS*_*WBVoff*_).

To remove motion artefacts from *WBV*_*off*_ and *WBV*_*on*_ epochs (Fratini et al., 2009a), a type II Chebyshev band-stop filter was applied at each stimulation frequency and its harmonics up to 450 Hz on the EMG spectra. This resulted in 30, 22, 18 ad 15 stop-band filters applied to epochs derived from WBV trials delivered at 15, 20, 25 and 30 Hz, respectively, following the calculation:

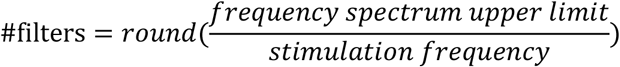

For each WBV trial, two *rRMS* vectors were computed on both artefact-free epochs (*WBV*_*off*_ and *WBV*_*on*_) (Fratini et al., 2009a) and used to calculate the relevant mean RMS values: *RMS*_*WBVoff∼*_ and *RMS*_*WBVon*_, respectively. To compare the values obtained during the different trials, a factor taking into account the proportion of power removed by the comb-notch filter was calculated (Lienhard et al., 2015):

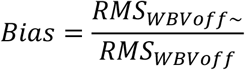

and was used to adjust *RMS*_*WBVon*_ values:

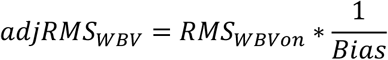

To evaluate the WBV-induced increment of muscular activation, *RMS*_*baseline*_ were subtracted from the *adjRMS*_*WBV*_ obtained for the WBV trials in the respective posture. These resulting values will be hereafter referred to as the *incrementRMS*_*WBV*_:

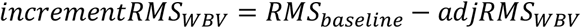

In total, eight values were retained for each subject and used for statistical analysis.

### Data processing and features extraction-Acceleration data

Baseline trials were not included in the following analyses: raw accelerations from WBV trials were analysed in Matlab ®R2019a (The Mathworks, Inc., Natick, MA). Accelerations were band-pass filtered between 10 and 100 Hz to remove gravity components and accommodation movements, usually confined between 0 and 5 Hz (Nowak et al., 2004; Prieto et al., 1996), and to retain only vibration-induced muscle displacements, located mostly at the stimulation frequency and its superior harmonics (Fratini et al., 2009a). Filtered epochs were then double integrated to estimate local displacement along the different axes (*disp*_*x*_, *disp*_*y*_, *disp*_*z*_) and the constant values introduced by each integration were removed with a 5 Hz high-pass filter. The total displacement recorded at each muscle level was estimated as:

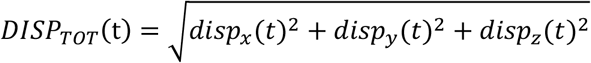

where *t* = 1,2, …, *N*, with *N* being the total number of samples.

To track the low-frequency mechanical muscle response to WBVs, a moving average of *DISP*_*TOT*_ (*MovAvgDISP*_*TOT*_) was calculated using a 250ms sliding window (Figure 1). To compare muscle displacement among different subjects, *MovAvgDISP*_*TOT*_ vectors were time-locked to the point where a 0.1 change in the slope was detected, which will be hereafter referred to as the vibration onset. *MovAvgDISP*_*TOT*_ from all participants were used for statistical analyses.

**Fig. 1.**
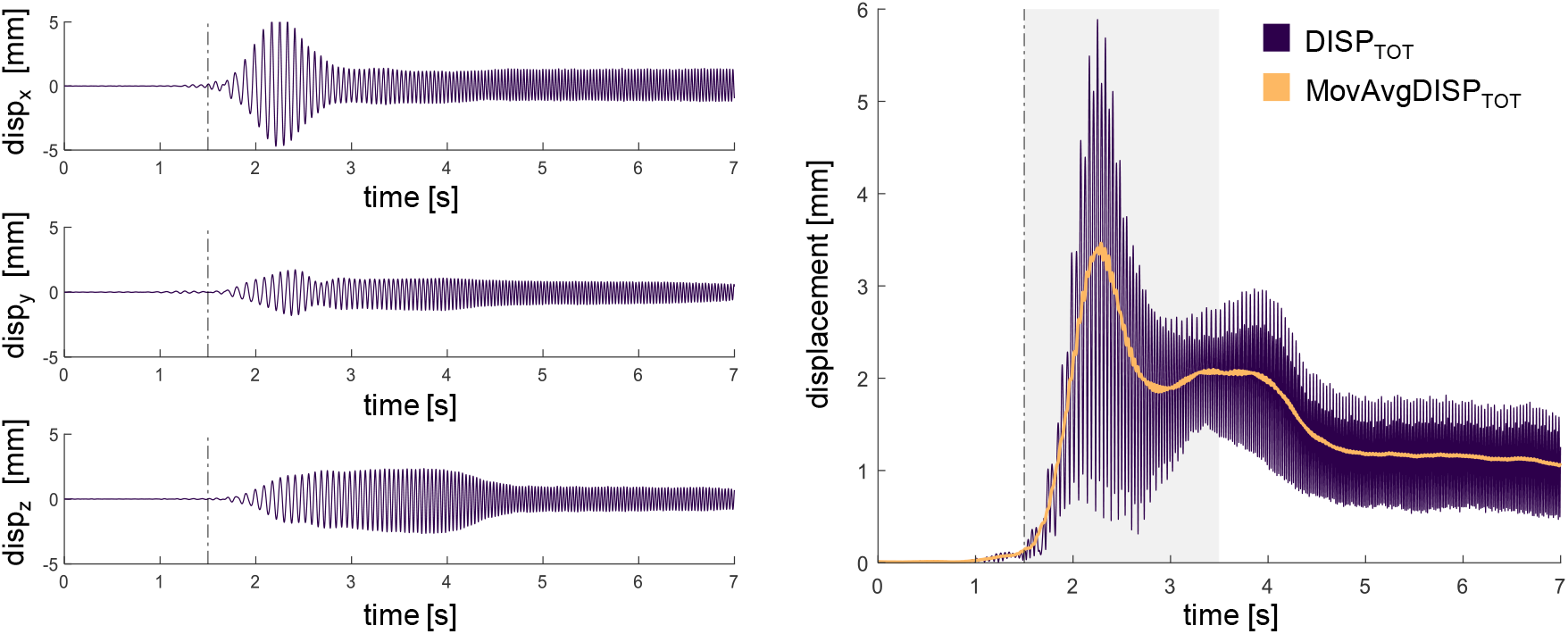
On the left side, muscle displacement obtained from double integration of the soft-tissue acceleration recorded at the GL site, HS_30_. Displacement along time is reported for the x, y and z axis. On the right, the GL total displacement obtained from the combination of the signals on the left (in purple): the moving average is depicted in orange. The vibration onset is indicated on the graphs by the vertical dashed line; the two-second interval used for the search of t_P_ is highlighted with a grey area

### Statistics

For each muscle, a cluster-based permutation test was used to compare the mechanical response of muscles over time (Maris, 2012; Maris and Oostenveld, 2007) for:

- *MovAvgDISP*_*TOT*_ between the two postures at different frequencies (four tests);
- *MovAvgDISP*_*TOT*_ between frequency pairs in HS and FF (twelve tests).

Time series comparisons were performed over the portion of the signals between the vibration onset and *t*_*A*_ to include both the peak and stabilization phase and because no effect was expected before the WBVs. 5000 permutations were used to build the random distribution against which the test statistic of the actual signal were compared. An alpha level of 0.05 was used to identify the significant clusters for each comparison (Gerber, 2020). To overcome the multiple comparison problem introduced by the number of comparisons run for each muscle, the cluster p values were further Bonferroni corrected.

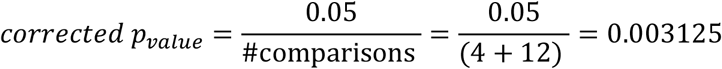

To test whether the electromyography activity increased significantly during WBVs, for each muscle, eight Wilcoxon signed rank tests (frequency (4) x posture (2)) compared the *incrementRMS*_*WBV*_ to a normal distribution with zero mean and unknown variance. Analysis were performed in Matlab ®R2019a.

For each analysed muscle, a two-way repeated measures Analysis of Variance (ANOVA) was conducted to examine the effect of stimulation frequency and subject posture on *incrementRMS*_*WBV*_ [frequency (4) x posture (2)], with Bonferroni corrections. Since muscle responses were investigated per se, outliers were removed from the dataset of the specific muscle after visual inspection of the data. Residuals were inspected and the approximate normal distribution of the data was confirmed by the Anderson-Darling test (Mohd Razali and Bee Wah, 2011). Mauchly’s test of sphericity was used to assess the sphericity of the data: when the latter was not met, a Greenhouse-Geisser correction was applied. Analysis were run in SPSS 23.0 (IBM Corp., Armonk, NY, USA) (Field, 2013).

## Results

All subjects were able to undergo WBV stimulations while holding the prescribed postures. The average ankle angles measured in Fore Feet were -9.35° ± 6.42 where a negative measure indicates a plantar flexion. When participants underwent the WBVs in Hack Squat, the average knee angle was 70.77° ± 4.43.

### Muscle dynamics analysis

Our results confirmed that the muscles dynamics differed significantly depending on the posture and frequency: overall, a larger displacement was observed in HS trials and at lower frequencies.

A characteristic mechanical peak was recorded shortly after the start of the stimulations in both postures, with the only exceptions of TA and SOL muscles when stimulated at 15 Hz. Moreover, although peaks varied among muscles, postures and frequencies, the average displacement showed a similar trend with a successive drop and a further stabilisation some seconds after the peak (Fig. 2).

**Fig. 2.**
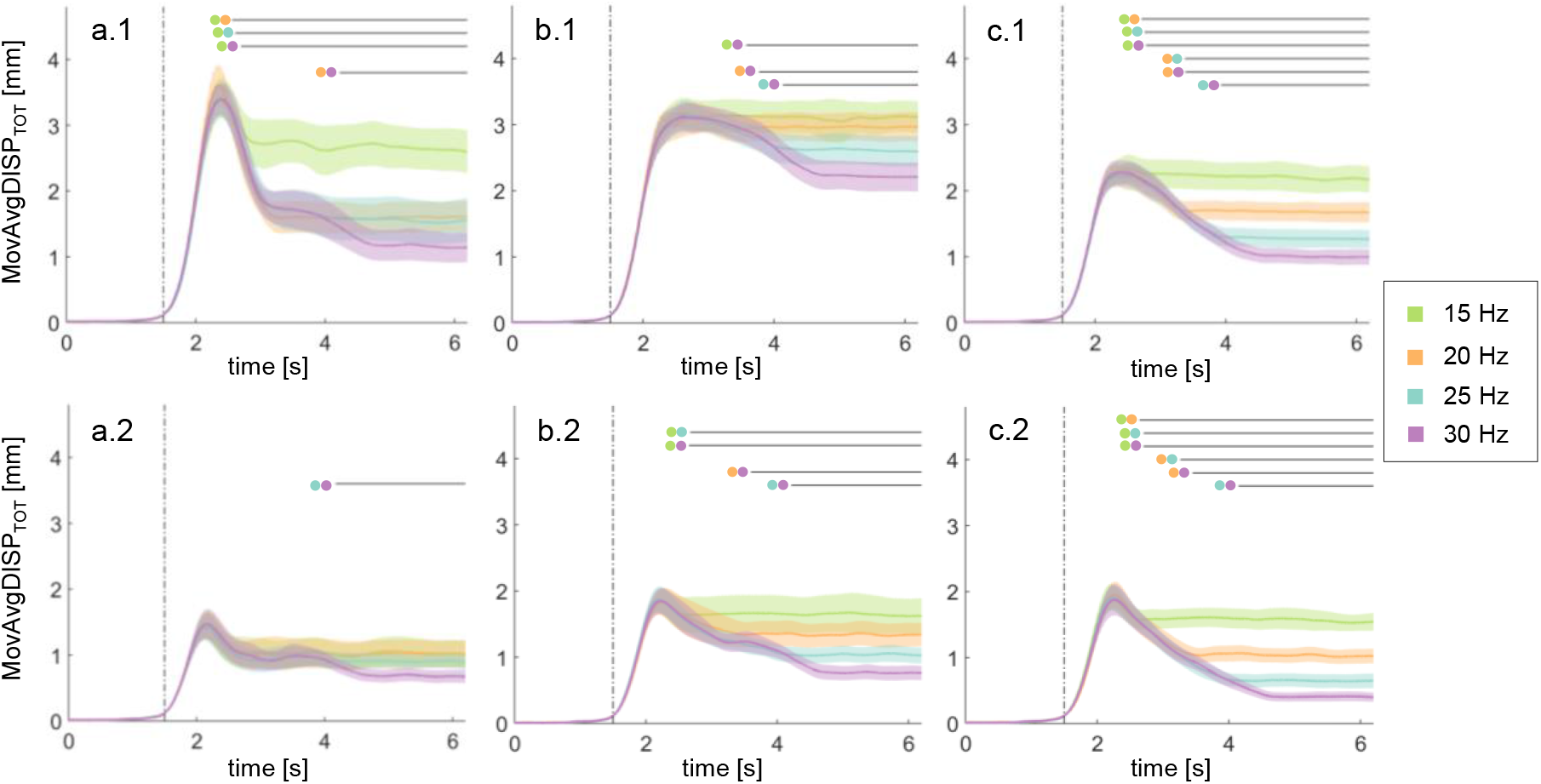
Moving average of the total displacement (mean +/- standard error) for each muscle (a=GL, b=SOL, c=TA) (N=25). The top row (.1) shows the mechanical responses while subjects underwent the WBVs in Hack Squat; the bottom row (.2) shows the responses while subjects were in Fore Feet. The results of the cluster-based permutation tests are indicated by the black lines (p<.003125) and the conditions considered for each comparison are listed via the colour-wise legend. The vertical dotted line represents the vibration onset

More in detail, although peak heights seemed stable across the frequency range, the drop changed significantly among muscles and postures (Figure 2). GL displacement after the peak, was significantly smaller at higher frequencies - at 20, 25 and 30 Hz rather than at 15 Hz (p=.0002) and at 30 Hz rather than at 20 (p=.0014) (Fig. 2, a.1) in HS. In FF, a similar trend was recorded: a smaller displacement was found at 30 Hz with respect to 25 Hz (p=.0004) (Fig. 2, a.2).

The average displacement recorded at the SOL site was smaller at 30 Hz than at 15 (p=.0006), 20 (p=.0002) and 25 Hz (p=.0014) while in HS (Fig. 2, b.1). Similarly, in FF, a smaller displacement was recorded at 30 Hz than at 15 (p=.0002), 20 (p=.001) and 25 Hz (p=.0002) and at 25 Hz than at 15 Hz (p=.0006) (Fig. 2, b.2).

The mechanical response of TA also confirmed the trend observed for the other two muscles. Its displacement was always smaller at higher frequencies in HS (*HS*_15_ < *HS*_20_, p=.0004; *HS*_25_ < *HS*_30_, p=.0006; for the other comparisons, p=.0002) (see Fig. 2, c.1) and in FF (p=.0002) (see Fig. 2, c.2).

### Muscle activity analysis

Normality was confirmed for the dependent variables in most of the conditions, for all three muscles. *incrementRMS*_*WBV*_ was not always normally distributed for TA, but the latter distributions were similarly skewed to those that met normality. Four subjects were removed from the *incrementRMS*_*WBV*_ dataset of GL and SOL, and three from that of TA, since represented outlier values. Distribution of *incrementRMS*_*WBV*_ values for the different muscles, posture and frequencies is depicted in Figure 3.

**Fig. 3.**
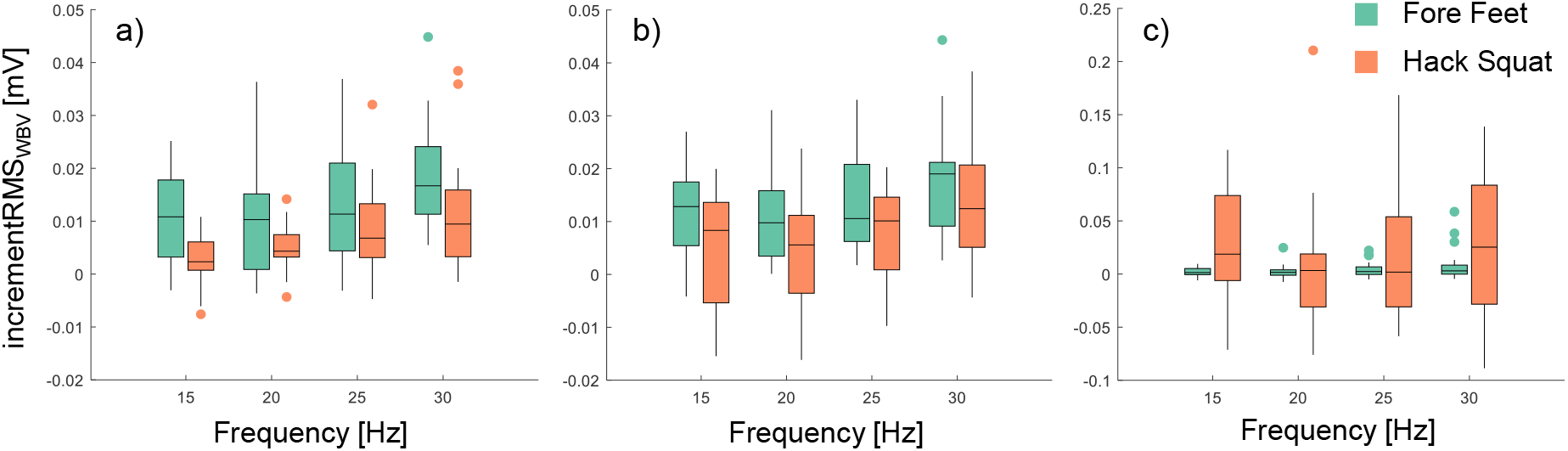
Box plots of incrementRMS_WBV_ values at different stimulation frequencies (15-30 Hz) of a=Gastrocnemius Lateralis (N=21), b=Soleus (N=21), and c=Tibialis Anterior (N=22) are shown. Different colours are used to distinguish between the muscle responses in hack squat (orange) and in fore feet (dark green) while the dots represent the outliers retained for the specific population. No significant interactions resulted from the ANOVAs. For significant main effects of stimulation frequency and subject posture refer back to the text. The figure was produced with Gramm (Morel, 2018)

A significant WBV-induced muscle activation (*incrementRMS*_*WBV*_) was observed in all conditions for the GL (see first row of Table 1) and in most of the conditions for the SOL, apart from *HS*_15_ (see second row of Table 1). Instead, the TA showed a significant response to WBVs only for 15 and 30 Hz and *FF*_25_ (see third row of Table 1).

**Table 1.**
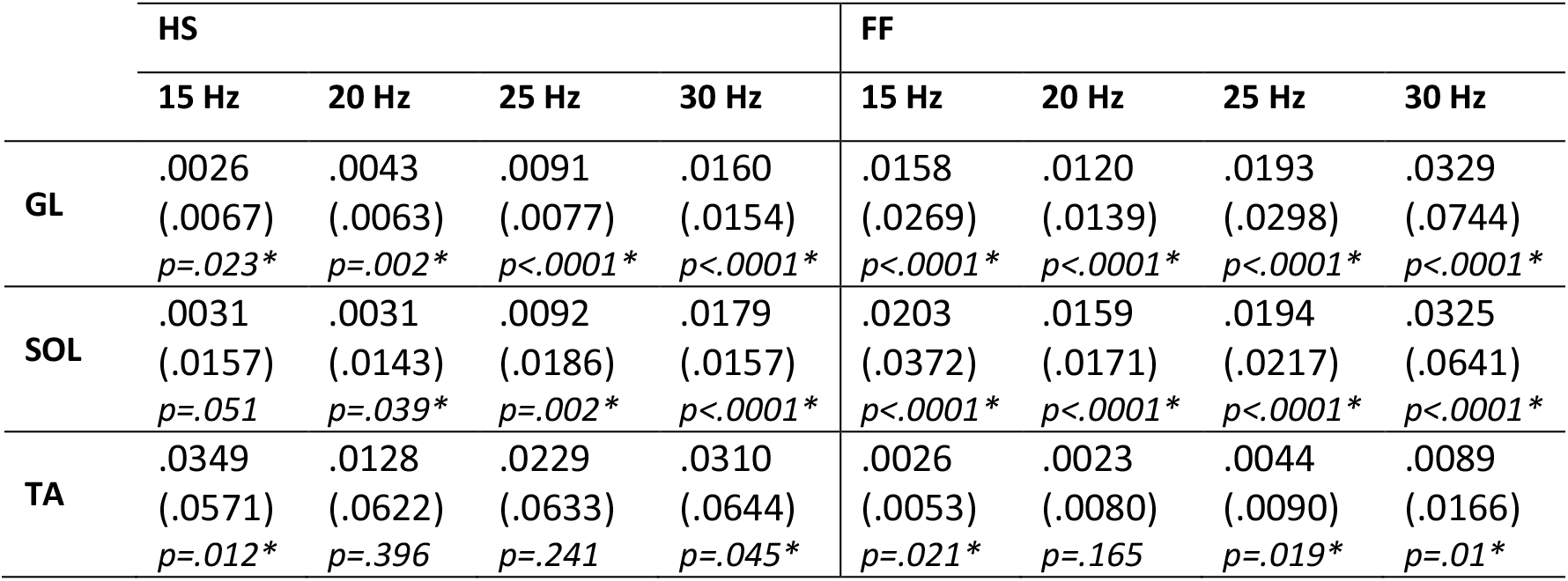
Results of the Wilcoxon signed rank tests used to test whether the WBV-induced increment of muscle activation (*incrementRMS*_*WBV*_) was significant. The mean (SD) of *incrementRMS*_*WBV*_ measured in each condition is reported, as well as the p-value of each test (N=25)

|  | HS |  |  |  | FF |  |  |  |
| --- | --- | --- | --- | --- | --- | --- | --- | --- |
|  | 15 Hz | 20 Hz | 25 Hz | 30 Hz | 15 Hz | 20 Hz | 25 Hz | 30 Hz |
| **GL** | .0026<br>(.0067)<br>$p=.023^*$ | .0043<br>(.0063)<br>$p=.002^*$ | .0091<br>(.0077)<br>$p<.0001^*$ | .0160<br>(.0154)<br>$p<.0001^*$ | .0158<br>(.0269)<br>$p<.0001^*$ | .0120<br>(.0139)<br>$p<.0001^*$ | .0193<br>(.0298)<br>$p<.0001^*$ | .0329<br>(.0744)<br>$p<.0001^*$ |
| **SOL** | .0031<br>(.0157)<br>$p=.051$ | .0031<br>(.0143)<br>$p=.039^*$ | .0092<br>(.0186)<br>$p=.002^*$ | .0179<br>(.0157)<br>$p<.0001^*$ | .0203<br>(.0372)<br>$p<.0001^*$ | .0159<br>(.0171)<br>$p<.0001^*$ | .0194<br>(.0217)<br>$p<.0001^*$ | .0325<br>(.0641)<br>$p<.0001^*$ |
| **TA** | .0349<br>(.0571)<br>$p=.012^*$ | .0128<br>(.0622)<br>$p=.396$ | .0229<br>(.0633)<br>$p=.241$ | .0310<br>(.0644)<br>$p=.045^*$ | .0026<br>(.0053)<br>$p=.021^*$ | .0023<br>(.0080)<br>$p=.165$ | .0044<br>(.0090)<br>$p=.019^*$ | .0089<br>(.0166)<br>$p=.01^*$ |

ANOVA analyses showed that although no significant interaction was found for the GL (N=21), main effect of stimulation frequency (F(3, 60) = 14.397, p< .0001) and subject posture were statistically significant (F(1, 20) = 15.433, p= .001). Specifically, GL-sEMG activity increased more in FF than in HS (p=.001) and 30 Hz was the stimulation frequency that evoked the highest muscular activation when compared to 15 Hz (p<.0001) and 20 Hz (p=.001). The WBV-induced increment of GL activation was also higher at 25 Hz than at 20 Hz (*p*=.02). Similarly, no significant interaction was found for the SOL (N=21) and a similar stimulation frequency (*F*(1.772, 35.434) = 12.982, p< .0001)) and subject posture (*F*(1, 20) = 6.357, p=.02)) main effects were found. The WBV-induced increment of SOL activity was higher in FF than in HS (p=.02) and 30 Hz was the stimulation frequency in which the highest sEMG increment was found when compared to 15 Hz (p=.002), 20 Hz (p=.004) and 25 Hz (p=.037). Moreover, a 25 Hz stimulation led to a higher muscle activation than 20 Hz (p=.009). No significant interaction nor main effect was instead found for the TA (N=22).

## Discussions

Although every muscle shows a peculiar response to WBVs - significantly determined by subject posture and stimulation frequency - a common mechanical pattern, never highlighted before, can be observed from our results. In response to vibratory stimulations, the extent of oscillations of muscles shows a rising phase, a peak oscillation and a subsequent drop, all of which completed within 4 to 5 seconds after the vibration onset, followed by a sustained stable response (plateau). Neither the stimulation amplitude nor the posture of participants varied during individual tests, hence a neuromuscular response must be responsible of this phenomenon. This interpretation is supported by the muscle tuning theory, which reports that soft-tissue oscillations arising in response of impact forces applied to the feet are dampened by an increase in muscle activation (Wakeling et al., 2003, 2002b; Wakeling and Nigg, 2001b). During WBVs, in fact, vibrations are transferred from the feet to the muscles via the body kinematic chain and produce soft-tissue compartment oscillations at the stimulation frequency, which in our case was in the range of the natural frequencies of calf muscles (Wakeling and Nigg, 2001a), and its harmonics (Burke and Schiller, 1976; Hagbarth et al., 1976; Hirayama et al., 1974; Homma et al., 1972; Martin and Park, 1997; Person and Kozhina, 1992, 1989). In light of the reported theory, it is therefore reasonable to assume that, if a resulting potential resonance is detected, muscle contraction is increased to avoid damage, creating the characteristic raising and falling curves observed in our recordings. This increase in muscle stiffness in fact, not only dampens the oscillations to a controlled level (observed plateau phase), but in turn shifts the natural frequency of the soft tissue toward higher values (Wakeling and Nigg, 2001a), and away from the WBV one. These dynamic analyses advance the understanding of muscles reaction to WBVs according to stimulation characteristics and, specifically, highlight that only after a period of time, which in this study is around 5 seconds, this reaction can completely settle. This may also explain why static exercises are found to be more effective than the dynamic ones (Abercromby et al., 2007a): during the first, muscles have time to tune to WBVs while during the latter the kinematic chain is continuously altered, leading to continuous changes in muscle contraction and sensitivity to vibrations (Burke et al., 1976).

The analysis of the physiological response of muscles to WBVs did instead highlighted specific combinations of posture/frequency able to produce maximal results. Recordings have been appropriately filtered for vibration-induced motion artefacts and therefore WBV-induced muscle activations were not overestimated (Fratini et al., 2014, 2009a; Romano et al., 2018). As expected, muscles behaved differently: GL activity was significantly enhanced in all WBV combinations, supporting the previously-demonstrated beneficial effects of WBV (Cardinale and Lim, 2003; Di Giminiani et al., 2013; Krol et al., 2011; Marin et al., 2009; Roelants et al., 2006), while for the SOL and the TA only specific combinations were able to significantly increase muscle activation. These findings highlight the importance of the selection of the most appropriate WBV parameters combinations, as it is clear that some settings are not effective in producing a greater activation of target muscles.

Among those combinations, undergoing WBV in Fore Feet was found to lead to a higher increase of GL and SOL sEMG activity rather than holding a Hack Squat position, confirming previous research findings (Ritzmann et al., 2013). Contracted muscles are in fact more responsive (Burke et al., 1976), and in our case this can be explained as GL and SOL are both plantar-flexors therefore more engaged in FF than HS (Carlsöö, 1961; Okada, 1972). WBVs delivered at 30 and 25 Hz triggered a greater activation in both muscles rather than 20 and 15 Hz, as similar findings reported (Di Giminiani et al., 2013) and supporting previous proposal of GL natural frequency residing between 25 and 30 Hz (Cesarelli et al., 2010). These conclusions are further confirmed by the observation of the permutation test results. Most differences were appreciable for the plateau phase, rather than the peak, where the displacement of GL and SOL soft-tissue compartments was significantly reduced at 30 Hz than at other WBV frequencies, further supporting the claim that this frequency is the one triggering the largest tuning effect. No specific effect was instead found for the TA. This results might be explained by (*i*) the stimulation frequencies used in this study that were limited to 30 Hz and not enough close to TA’s natural frequency, which ranges up to 50 Hz (Wakeling et al., 2002a); (*ii*) the selected postures that did not lead to an appropriate level of TA engagement, limiting its response to WBVs (Burke et al., 1976).

Combining the above, it can be inferred that 30 Hz-Fore Feet might be the best combination of stimulation frequency and subject posture when aiming to effectively enhance both GL and SOL muscular responses. For the explored combinations, instead, the TA muscle showed that WBVs elicit muscular activity but did not allow to identify any combination producing a significantly higher response. Therefore, a wider range of frequencies and postures should be explored.

## Conclusions

Our results highlighted that WBVs via side-alternating platforms can be optimised for calf muscle training, if aimed at achieving the maximal level of elicited response, and in particular:

- In order to allow muscles to produce a stable contraction, training programmes should only include static exercise on a vibrating platform or, at least, participants should hold the same posture for a minimum of five seconds.
- Calf muscles produce maximal response if participants are standing on the fore feet during stimulations.
- Vibration frequencies in the range of 25-30Hz should be used during WBV training sessions for calf muscles.

The systematic targeting of the plantar flexors via proper selection of stimulation frequency and subject posture represents an opportunity to maximise the ankle strategy outcome and the possibly improve postural control mechanisms.

Future studies aimed at investigating both the postural acute and chronic effect on calf muscle-training via mean of WBVs are desirable to further confirm and expand the findings of this study.

## Abbreviations

ANOVA: Analysis of variance
sEMG: (surface) Electromyography
FF: Fore feet
GL: Gastrocnemius lateralis
HS: Hack squat
SOL: Soleus
RMS: Root mean square
TA: Tibialis anterior
TVR: Tonic vibration reflex
WBV: Whole body vibration

## Declarations

### Availability of data and material

Since sharing data in an open-access repository was not included in our participant’s consent and therefore compromises our ethical standards, data are only available on request from the corresponding author.

### Code availability

The code used for the analyses of the data will be shared upon request to the corresponding author

### Author’s contributions

Conception and design: I. R., T. B., P. B. and A. F. Data acquisition and analysis: I. R. Interpretation: I. R. and A. F. Drafting manuscript: I. R., T. B., P.B. and A. F.

### Ethics approval

The study was carried out according to the Declaration of Helsinki (2013) and was approved by the University Research Ethics Committee at Aston University (reference number: 1439).

### Consent to participate

All participants provided informed consent before participating.

### Consent for publication

All co-authors were aware of the publication of this study

## References

Abercromby, A., Amonette, W., Layne, C., Mcfarlin, B., Hinman, M., Paloski, W., 2007a. Variation in neuromuscular responses during acute whole-body vibration exercise. Med. Sci. Sports Exerc. 39, 1642–1650. https://doi.org/10.1249/mss.0b013e318093f551

Abercromby, A., Amonette, W., Layne, C., Mcfarlin, B., Hinman, M., Paloski, W., 2007b. Vibration exposure and biodynamic responses during whole-body vibration training. Med. Sci. Sports Exerc. 39, 1794–1800. https://doi.org/10.1249/mss.0b013e3181238a0f

Alam, M.M., Khan, A.A., Farooq, M., 2018. Effect of whole-body vibration on neuromuscular performance: A literature review. Work 59, 571–583. https://doi.org/10.3233/WOR-182699

AX3 OMGUI Configuration and Analysis Tool, 38, GitHub, 2015.

Bautmans, I., Van Hees, E., Lemper, J.C., Mets, T., 2005. The feasibility of whole body vibration in institutionalised elderly persons and its influence on muscle performance, balance and mobility: A randomised controlled trial [ISRCTN62535013]. BMC Geriatr. 5, 1–8. https://doi.org/10.1186/1471-2318-5-17

Bogaerts, A., Verschueren, S., Delecluse, C., Claessens, A.L., Boonen, S., 2007. Effects of whole body vibration training on postural control in older individuals: A 1 year randomized controlled trial. Gait Posture 26, 309–316. https://doi.org/10.1016/j.gaitpost.2006.09.078

Bosco, C., Colli, R., Introini, E., Cardinale, M., Tsarpela, O., Madella, a, Tihanyi, J., Viru, a, 1999. Adaptive responses of human skeletal muscle to vibration exposure. Clin. Physiol. 19, 183–187. https://doi.org/10.1046/j.1365-2281.1999.00155.x

Burke, D., Hagbarth, K.E., Lofstedt, L., Wallin, B.G., 1976. The responses of human muscle spindle endings to vibration during isometric contraction. J. Physiol. 261, 695–711. https://doi.org/10.1113/jphysiol.1976.sp011581

Burke, D., Schiller, H.H., 1976. Discharge pattern of single motor units in the tonic vibration reflex of human triceps surae. J. Neurol. Neurosurg. Psychiatry 39, 729–741. https://doi.org/10.1136/jnnp.39.8.729

Cardinale, M., Lim, J., 2003. Electromyography activity of vastus lateralis muscle during whole-body vibrations of different frequencies. J. Strength Cond. Res. 17, 621–624. https://doi.org/10.1519/1533-4287(2003)017<0621:EAOVLM>2.0.CO;2

Carlsöö, S., 1961. The static muscle load in different work positions: An electromyographic study. Ergonomics 4, 193–211. https://doi.org/10.1080/00140136108930520

Cesarelli, M., Fratini, A., Bifulco, P., La Gatta, A., Romano, M., Pasquariello, G., 2010. Analysis and modelling of muscles motion during whole body vibration. EURASIP J. Adv. Signal Process. 2010, 26–28. https://doi.org/10.1155/2010/972353

Delecluse, C., Roelants, M., Verschueren, S., 2003. Strength increase after whole-body vibration compared with resistance training. Med. Sci. Sports Exerc. 35, 1033–1041. https://doi.org/10.1249/01.MSS.0000069752.96438.B0

Di Giminiani, R., Masedu, F., Tihanyi, J., Scrimaglio, R., Valenti, M., 2013. The interaction between body position and vibration frequency on acute response to whole body vibration. J. Electromyogr. Kinesiol. 23, 245–251. https://doi.org/10.1016/j.jelekin.2012.08.018

Dolny, D.G., Reyes, F.C.G., 2008. Whole body vibration exercise: Training and benefits. Curr. Sports Med. Rep. 7, 152–157. https://doi.org/10.1097/01.CSMR.0000319708.18052.a1

Fagnani, F., Giombini, A., Di Cesare, A., Pigozzi, F., Di Salvo, V., 2006. The effects of a whole-body vibration program on muscle performance and flexibility in female athletes. Am. J. Phys. Med. Rehabil. 85, 956–962. https://doi.org/10.1097/01.phm.0000247652.94486.92

Field, A., 2013. Discovering Statistics using IBM SPSS statistics, 4th ed. SAGE Publication Ltd.

Fitzpatrick, R.C., Gorman, R.B., Burke, D., Gandevia, S.C., 1992. Postural proprioceptive reflexes in standing human subjects: bandwidth of response and transmission characteristics. J. Physiol. 458, 69–83.

Fratini, A., Bifulco, P., Romano, M., Clemente, F., Cesarelli, M., 2014. Simulation of surface EMG for the analysis of muscle activity during whole body vibratory stimulation. Comput. Methods Programs Biomed. 113, 314–322. https://doi.org/10.1016/j.cmpb.2013.10.009

Fratini, A., Cesarelli, M., Bifulco, P., Romano, M., 2009a. Relevance of motion artifact in electromyography recordings during vibration treatment. J. Electromyogr. Kinesiol. 19, 710–718. https://doi.org/10.1016/j.jelekin.2008.04.005

Fratini, A., La Gatta, A., Cesarelli, M., Bifulco, P., 2009b. Whole Body Vibration training: analysis and characterization. 2009 9th Int. Conf. Inf. Technol. Appl. Biomed. 1–4. https://doi.org/10.1109/ITAB.2009.5394317

Gatev, P., Thomas, S., Kepple, T., Hallett, M., 1999. Feedforward ankle strategy of balance during quiet stance in adults. J. Physiol. 514, 915–928. https://doi.org/10.1111/j.1469-7793.1999.915ad.x

Gerber, E.M., 2020. permutest [WWW Document]. URL https://uk.mathworks.com/matlabcentral/fileexchange/71737-permutest (accessed 5.20.20).

Granit, R., Steg, G., 1956. Tonic and Phasic Ventral Horn Cells Differentiated by Post-Tetanic Potentiation in Cat Extensors. Acta Physiol Scand 37, 114–126.

Gross, T.S., Nelson, R.C., 1988. The shock attenuation role of the ankle during landing from a vertical jump. Med. Sci. Sports Exerc. 20, 506–514.

Hagbarth, K.E., Hellsing, G., Löfstedt, L., 1976. TVR and vibration-induced timing of motor impulses in the human jaw elevator muscles. J. Neurol. Neurosurg. Psychiatry 39, 719–728. https://doi.org/10.1136/jnnp.39.8.719

Harazin, B., Grzesik, J., 1998. The transmission of vertical whole-body vibration to the body segments of standing subjects. J. Sound Vib. 215, 775–787. https://doi.org/10.1006/jsvi.1998.1675

Hermens, H.J., Bart, F., Catherine, D.-K., Gunter, R., 2000. Development of recommendations for SEMG sensors and sensor placement procedures. J. Electromyogr. Kinesiol. 10, 361–374.

Hirayama, K., Homma, S., Mizote, M., Nakajima, Y., Watanabe, S., 1974. Separation of the contributions of voluntary and vibratory activation of motor units in man by cross-correlograms. Jpn. J. Physiol. 24, 293–304.

Homma, S., Kanda, K., Watanabe, S., 1972. Preferred spike intervals in the vibration reflex. Jpn. J. Physiol. 22, 421–432.

Horak, F.B., Sharon, S.M., Shumway-Cook, A., 1997. Postural Pertubations: New Insights for Treatment of Balance Disorders. Phys. Ther. 77, 517–533.

Krol, P., Piecha, M., Slomka, K., Sobota, G., Polak, A., Juras, G., 2011. The effect of whole-body vibration frequency and amplitude on the myoelectric activity of vastus medialis and vastus lateralis. J. Sport. Sci. Med. 10, 169–174.

Lafortune, M.A., Lake, M.J., 1996. Dominant Role of Interface Over Knee Angle for Cushioning Impact Loading and Regulating Initial Leg Stiffness. J. Biomech. 29, 1523–1529.

Lafortune, M.A., Lake, M.J., Hennig, E.M., 1996. Differential shock transmission response of the human body to impact severity and lower limb posture. J. Biomech. 29, 1531–1537. https://doi.org/10.1016/S0021-9290(96)80004-2

Lam, F.M.H., Lau, R.W.K., Chung, R.C.K., Pang, M.Y.C., 2012. The effect of whole body vibration on balance, mobility and falls in older adults: A systematic review and meta-analysis. Maturitas 72, 206–213. https://doi.org/10.1016/j.maturitas.2012.04.009

Lienhard, K., Cabasson, A., Meste, O., Colson, S.S., 2015. Comparison of sEMG processing methods during whole-body vibration exercise. J. Electromyogr. Kinesiol. 25, 833–840. https://doi.org/10.1016/j.jelekin.2015.10.005

Lienhard, K., Cabasson, A., Meste, O., Colson, S.S., 2014. Determination of the optimal parameters maximizing muscle activity of the lower limbs during vertical synchronous whole-body vibration. Eur. J. Appl. Physiol. 114, 1493–1501. https://doi.org/10.1007/s00421-014-2874-1

Mahieu, N.N., Witvrouw, E., Van De Voorde, D., Michilsens, D., Arbyn, V., Van Den Broecke, W., 2006. Improving strength and postural control in young skiers: Whole-body vibration versus equivalent resistance training. J. Athl. Train. 41, 286–293.

Marin, P., Bunker, D., Rhea, M., Ayllon, F., 2009. Neuromuscular Activity During Whole Body Vibration of Different Amplitudes and Footwear Conditions: Implications for Prescription of Vibratory Stimulation. J. Strength Cond. Res. 23, 2311–2316.

Maris, E., 2012. Statistical testing in electrophysiological studies. Psychophysiology 49, 549–565. https://doi.org/10.1111/j.1469-8986.2011.01320.x

Maris, E., Oostenveld, R., 2007. Nonparametric statistical testing of EEG- and MEG-data. J. Neurosci. Methods 164, 177–190. https://doi.org/10.1016/j.jneumeth.2007.03.024

Martin, B.J., Park, H.S., 1997. Analysis of the tonic vibration reflex: Influence of vibration variables on motor unit synchronization and fatigue. Eur. J. Appl. Physiol. 75, 504–511. https://doi.org/10.1007/s004210050196

Mohd Razali, N., Bee Wah, Y., 2011. Power comparisons of Shapiro-Wilk, Kolmogorov-Smirnov, Lilliefors and Anderson-Darling tests. J. Stat. Model. Anal. 2, 21–33.

Morel, P., 2018. Gramm: grammar of graphics plotting in Matlab. J. Open Source Softw. 3, 568. https://doi.org/10.21105/joss.00568

Nowak, D.A., Rosenkranz, K., Hermsdörfer, J., Rothwell, J., 2004. Memory for fingertip forces: Passive hand muscle vibration interferes with predictive grip force scaling. Exp. Brain Res. 156, 444–450. https://doi.org/10.1007/s00221-003-1801-1

Okada, M., 1972. An electromyographic estimation of the relative muscular load in different human postures. J. Hum. Ergol. (Tokyo). 1, 75–93. https://doi.org/10.11183/jhe1972.1.75

Osawa, Y., Oguma, Y., Ishii, N., 2013. The effects of whole-body vibration on muscle strength and power: A meta-analysis. J. Musculoskelet. Neuronal Interact. 13, 342–352.

Person, R., Kozhina, G., 1992. Tonic vibration reflex of human limb muscles: Discharge pattern of motor units. J. Electromyogr. Kinesiol. 2, 1–9. https://doi.org/10.1016/1050-6411(92)90002-Z

Person, R.S., Kozhina, G.V., 1989. Study of Firing Pattern in Human Soleus Motor Units in Tonic Vibration Reflex. Neurophysiology 21, 540–546.

Peterka, R.J., 2002. Sensorimotor integration in human postural control. J. Neurophysiol. 88, 1097–1118. https://doi.org/10.1152/jn.2002.88.3.1097

Pollock, R.D., Woledge, R.C., Martin, F.C., Newham, D.J., 2012. Effects of whole body vibration on motor unit recruitment and threshold. J. Appl. Physiol. 112, 388–395. https://doi.org/10.1152/japplphysiol.01223.2010

Pollock, R.D., Woledge, R.C., Mills, K.R., Martin, F.C., Newham, D.J., 2010. Muscle activity and acceleration during whole body vibration: Effect of frequency and amplitude. Clin. Biomech. 25, 840–846. https://doi.org/10.1016/j.clinbiomech.2010.05.004

Prieto, T.E., Myklebust, J.B., Hoffmann, R.G., Lovett, E.G., Myklebust, B.M., 1996. Measures of postural steadiness: Differences between healthy young and elderly adults. IEEE Trans. Biomed. Eng. 43, 956–966. https://doi.org/10.1109/10.532130

Rittweger, J., 2010. Vibration as an exercise modality: How it may work, and what its potential might be. Eur. J. Appl. Physiol. 108, 877–904. https://doi.org/10.1007/s00421-009-1303-3

Ritzmann, R., Gollhofer, A., Kramer, A., 2013. The influence of vibration type, frequency, body position and additional load on the neuromuscular activity during whole body vibration. Eur. J. Appl. Physiol. 113, 1–11. https://doi.org/10.1007/s00421-012-2402-0

Ritzmann, R., Kramer, A., Bernhardt, S., Gollhofer, A., 2014. Whole body vibration training - Improving balance control and muscle endurance. PLoS One 9. https://doi.org/10.1371/journal.pone.0089905

Ritzmann, R., Kramer, A., Gruber, M., Gollhofer, A., Taube, W., 2010. EMG activity during whole body vibration: Motion artifacts or stretch reflexes? Eur. J. Appl. Physiol. 110, 143–151. https://doi.org/10.1007/s00421-010-1483-x

Roelants, M., Verschueren, S.M.P., Delecluse, C., Levin, O., Stijnen, V., 2006. Whole-body-vibration-induced increase in leg muscle activity during different squat exercises. J. Strength Cond. Res. 20, 124–129. https://doi.org/10.1519/R-16674.1

Rogan, S., Hilfiker, R., Herren, K., Radlinger, L., De Bruin, E.D., 2011. Effects of whole-body vibration on postural control in elderly: A systematic review and meta-analysis. BMC Geriatr. 11. https://doi.org/10.1186/1471-2318-11-72

Romano, M., Fratini, A., Gargiulo, G.D., Cesarelli, M., Iuppariello, L., Bifulco, P., 2018. On the Power Spectrum of Motor Unit Action Potential Trains Synchronized With Mechanical Vibration. IEEE Trans. Neural Syst. Rehabil. Eng. 26, 646–653.

Saquetto, M., Carvalho, V., Silva, C., Conceição, C., Gomes-Neto, M., 2015. The effects of whole body vibration on mobility and balance in children with cerebral palsy: A systematic review with meta-analysis. J. Musculoskelet. Neuronal Interact. 15, 137–144.

Torvinen, S., Kannus, P., Sievänen, H., Järvinen, T.A.H., Pasanen, M., Kontulainen, S., Järvinen, T.L.N., Järvinen, M., Oja, P., Vuori, I., 2002a. Effect of a vibration exposure on muscular performance and body balance. Randomized cross-over study. Clin. Physiol. Funct. Imaging 22, 145–152. https://doi.org/10.1046/j.1365-2281.2002.00410.x

Torvinen, S., Kannus, P., Sievanen, H., Vuori, I., Paakkala, T., Ilkka, V., Al., E., 2003. Effect of 8-month vertical whole body vibration on bone, muscle performance, and body balance: a randomized controlled study. J. Bone Miner. Res. 18, 876–884.

Torvinen, S., Sievanen, H., Jarvinen, T.A.H., Pasanen, M., Kontulainen, S., Kannus, P., 2002b. Effect of 4-min vertical whole body vibration on muscle performance and body balance: A randomized cross-over study. Int. J. Sports Med. 23, 374–379. https://doi.org/10.1055/s-2002-33148

Turbanski, S., Haas, C.T., Schmidtbleicher, D., Friedrich, A., Duisberg, P., 2005. Effects of random whole-body vibration on postural control in Parkinson’s disease. Res. Sport. Med. 13, 243–256. https://doi.org/10.1080/15438620500222588

van Nes, I.;, Geurts, A.;, Hendricks, H.T.., Duysens, J., 2004. Short-Term Effects of Whole-Body Vibration on Postural Control in Unilateral Chronic Stroke Patients: Preliminary Evidence. Am. J. Phys. Med. Rehabil. 83, 867–873.

Wakeling, J., Liphardt, A.M., Nigg, B., 2003. Muscle activity reduces soft-tissue resonance at heel-strike during walking. J. Biomech. 36, 1761–1769. https://doi.org/10.1016/S0021-9290(03)00216-1

Wakeling, J., Nigg, B., 2001a. Modification of soft tissue vibrations in the leg by muscular activity. J Appl Physiol 90, 412–420. https://doi.org/10.1152/jappl.2001.90.2.412

Wakeling, J., Nigg, B., 2001b. Soft-tissue vibrations in the quadriceps measured with skin mounted transducers. J. Biomech. 34, 539–543.

Wakeling, J., Nigg, B., Rozitis, A., 2002a. Muscle activity damps the soft tissue resonance that occurs in response to pulsed and continuous vibrations. J. Appl. Physiol. 93, 1093–1103. https://doi.org/10.1152/japplphysiol.00142.2002

Wakeling, J., Pascual, S., Nigg, B., 2002b. Altering muscle activity in the lower extremities by running with different shoes. Med. Sci. Sport. Exerc. 34, 1529–1532. https://doi.org/10.1249/01.MSS.0000027714.70099.08

Wakeling, J., Von Tscharner, V., Nigg, B., Stergiou, P., 2001. Muscle activity in the leg is tuned in response to ground reaction forces. J. Appl. Physiol. 91, 1307–1317.

Winter, D.A.., 1995. Human balance and posture control during standing and walking. Gait Posture 3, 193–214. https://doi.org/10.1016/0014-5793(86)80927-9

Wyon, M., Guinan, D., Hawkey, A., 2010. Whole-Body vibration training increases vertical jump height in a dance population. J. Strength Cond. Res. 24, 866–870.

Yang, X., Wang, P., Liu, C., He, C., Reinhardt, J.D., 2015. The effect of whole body vibration on balance, gait performance and mobility in people with stroke: A systematic review and meta-analysis. Clin. Rehabil. 29, 627–638. https://doi.org/10.1177/0269215514552829

